# Local variations in L/M ratio influence the detection and color naming of small spots

**DOI:** 10.1101/2025.02.19.639104

**Authors:** Maxwell J. Greene, Vimal P. Pandiyan, Ramkumar Sabesan, William S. Tuten

## Abstract

The distribution of long (L), middle (M), and short (S) wavelength sensitive cones in the retina determines how different frequencies of incident light are sampled across space and has been hypothesized to influence spatial and color vision. We examined how the detection and color naming of small, short-duration increment stimuli (λ = 543 or 680 nm) depend on the local spectral topography of the cone mosaic. Stimuli were corrected for optical aberrations by an adaptive optics system and targeted to locations in the parafovea where cone spectral types were known. We found that sensitivity to 680 nm light, normalized by sensitivity to 543 nm light, grew with the proportion of L cones at the stimulated locus, though intra- and intersubject variability was considerable. A similar trend was derived from a simple model of the achromatic (L+M) pathway suggesting that small spot detection mainly relies on a non-opponent mechanism. Most stimuli were called achromatic, with red and green responses becoming more common as stimulus intensity increased and as the local L/M ratio became more symmetric. The proximity of S cones to the stimulated region did not influence the likelihood of eliciting a chromatic percept. Our detection data confirm earlier reports that small spot psychophysics can reveal information about local cone topography, and our color naming findings suggest that chromatic sensitivity may improve when the L/M ratio approaches unity.

## INTRODUCTION

The relative numbers of long (L) and middle (M) wavelength sensitive cones in the eye can vary substantially between human trichromats. The existence of such diversity at the first stage of retinal processing was first supposed based on intersubject differences in luminous efficiencies (de Vries, 1946, 1949), and later confirmed by molecular analyses (Hagstrom et al., 1998, 2000), electroretinographic recordings (Brainard et al., 1999; Carroll et al., 2002), and *in vivo* adaptive optics imaging (Hofer, Carroll, et al., 2005; Pandiyan et al., 2022; Roorda & Williams, 1999; Zhang et al., 2019). In addition to variations in L/M ratio between subjects, the spatial distribution of L and M cones within an individual primate retina is highly non-uniform, resulting in pockets of the mosaic where the local proportion of L cones deviates from the overall average (Hofer, Carroll, et al., 2005; Mollon & Bowmaker, 1992; Roorda et al., 2001).

How L/M ratio variability within and between individuals affects vision is not yet fully understood. This is because the topography of the human cone mosaic, and its effect on downstream visual processing, have historically been unamenable to direct study. Adaptive optics-based technology is well suited to advance our understanding in this area, since it enables the spectral classification of single cones, and in turn the examination of how perception depends on stimulating cones of known spectral type (Fong et al., 2025; Greene et al., 2024; Sabesan et al., 2016; Schmidt, Boehm, et al., 2018; Schmidt et al., 2019).

The retinal L/M cone ratio has been thought to influence various aspects of visual performance. Consider an observer whose photoreceptor mosaic contains a higher-than-average number of L cones (i.e., a high L/M ratio). Two hypotheses can be made about their visual system. First, compared to a typical observer with an L/M ratio of ∼2:1 (Carroll et al., 2002), they should be relatively more sensitive to long-wavelength light. This hypothesis is broadly supported by sensitivity measurements obtained with both large-field stimuli (Kremers et al., 2000; Rushton & Baker, 1964) and small flashes (Cicerone & Nerger, 1989; Vimal et al., 1989), although inferring cone numbers solely from threshold data requires making assumptions about the properties of the receptors themselves (e.g., spectral sensitivity, optical density; cf. Bieber et al., 1998; He et al., 2021), as well as how their signals are processed by post-receptoral circuits (Stockman et al., 2008).

Second, a more homogeneous cone mosaic may be detrimental to color vision. Extracting spectral information from the retinal image depends on circuits that compare signals from more than one cone class. If one cone type outnumbers the other, the fidelity of cone-opponent signals may be limited by noise in the minority receptor class. Previous work suggests the detectability and discriminability of low-frequency (≤ 2 cpd) red-green gratings is reduced in retinas with presumed asymmetric L/M ratios (Gunther & Dobkins, 2002; Hood et al., 2006). Likewise, the ability to resolve finer-grained spatiochromatic patterns also appears to be constrained by the spectral makeup of the photoreceptor layer (Danilova et al., 2013; Neitz et al., 2020).

In the present study, we examine both hypotheses about the local L/M ratio by measuring the detectability and color appearance of small, brief, narrowband (λ = 543 and 680 nm) increments delivered via adaptive optics scanning light ophthalmoscopy (AOSLO; Roorda et al., 2002). Although S cones are insensitive to the wavelengths we used (Stockman et al., 1999), we also evaluated whether the proximity of the stimulus to S cones influenced small-spot color perception, as has been suggested by earlier studies (Brainard et al., 2008). Advantages of the investigation described here over older ones concerned with similar questions are (1) objective cone spectral classifications via optoretinography were available for each subject (Pandiyan et al., 2022), and (2) the use of AOSLO enabled the controlled delivery of stimuli to regions that varied in local L/M ratio. Our threshold data and modeling suggest that the primary mechanism mediating detection of our stimuli is one that sums L and M cones with a weight that does not depend on the local arrangement of L and M cones, although our color appearance findings suggest a secondary contribution from chromatic mechanisms. Furthermore, after equating for detectability, subjects were more likely to categorize the stimuli as chromatic when they were sampled by similar numbers of L and M cones, whereas stimuli landing on more homogenous retinal loci tended to yield achromatic percepts. Proximity to S cones was not a significant predictor of a stimulus being perceived as chromatic.

## METHODS

### Subjects

Two authors of the present study volunteered as experimental observers (subjects 10001R and 20217R). Prior to study enrollment, subjects gave informed written consent. All study procedures adhered to the tenets of the 2008 Declaration of Helsinki and were approved by institutional review boards at the University of California, Berkeley and the University of Washington. Both subjects were males (ages 40 and 27 years) with normal color vision as assessed by a Neitz anomaloscope and/or pseudoisochromatic plates viewed under standard illuminant C. Both participants have extensive experience as adaptive optics psychophysics subjects. The subjects’ cone spectral identities were previously determined by optoretinography at the University of Washington, a technique based on adaptive optics optical coherence tomography (AO-OCT; Pandiyan et al., 2020). Cone classification with AO-OCT is faster and more sensitive than AO-densitometry (Pandiyan et al., 2022; Zhang et al., 2019). Cone types were known within a 1.37 x 1.1 deg patch of retina at 2 deg temporal eccentricity for subject 10001R and within a 0.86 x 0.93 deg patch of retina at 2.5 deg temporal eccentricity for subject 20217R. Subregions of these cone-classified mosaics are shown in **Figure 1A**. Within his classified region, subject 10001R’s L/M ratio is 1.47, while subject 20217R’s L/M ratio is 3.66. Because the cone classification procedure required travel from Berkeley to Seattle, it was not practical to recruit naïve subjects for this study. To minimize the possibility of observer bias influencing the subjective color reports, we designed the psychophysical paradigm to ensure that subjects could not predict the wavelength, intensity, or locus of stimulus delivery on the basis of the preceding trial (see Methods section entitled ***“Psychophysics”*** below).

**Figure 1.**
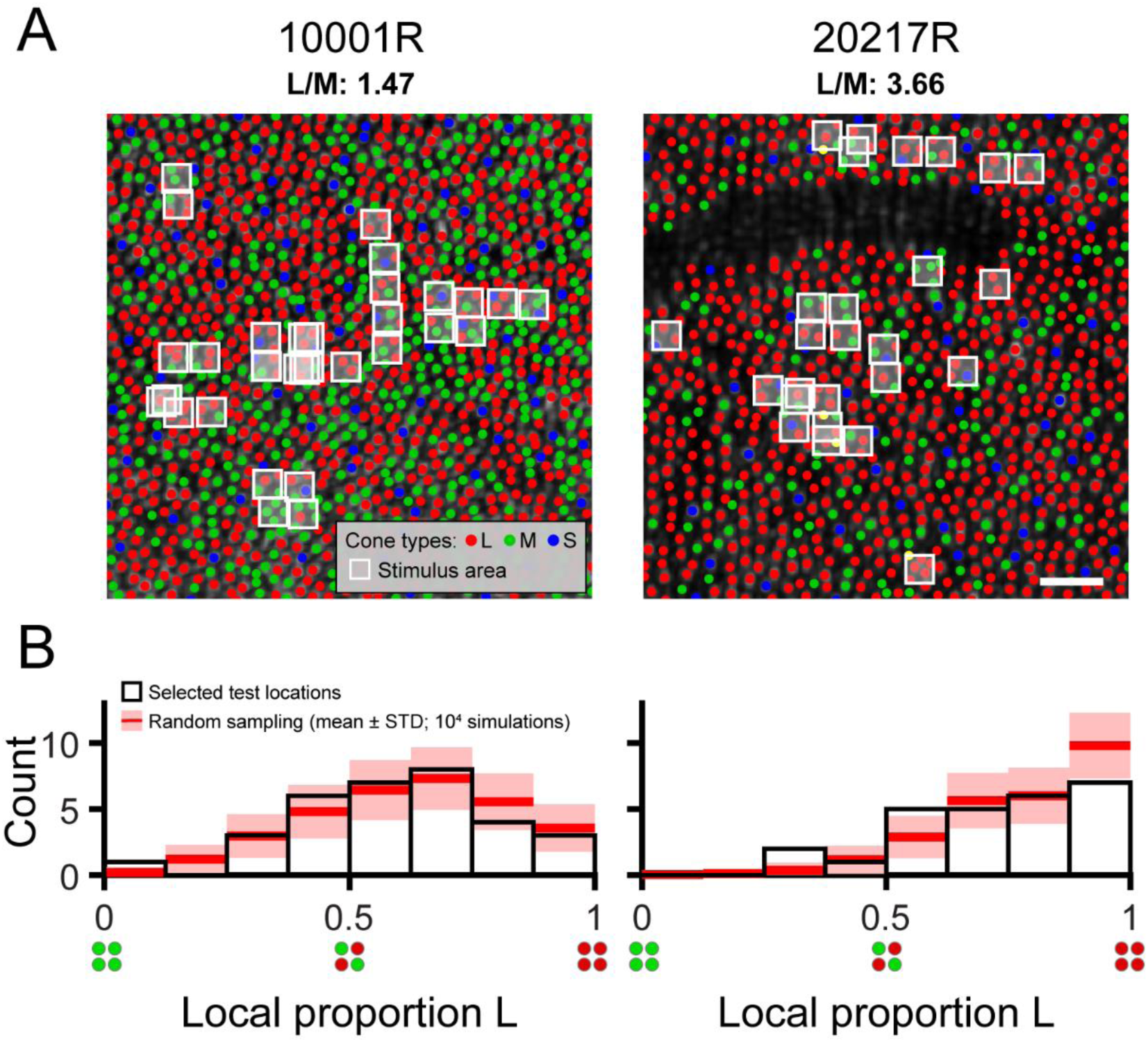
Local variation in cone spectral topography in the trichromatic retina. (**A**) Labeled cone mosaic images acquired from the two participants in this study (0.65° x 0.65° field of view). L, M, and S cones are labeled using red, green, and blue markers, respectively. The mean delivery coordinate for each tested location is marked by a white box whose size represents the nominal stimulus area on the retina (2.25 x 2.25 arcmin). The white scale bar corresponds to a visual angle of 5 arcmin. The dark unlabeled region in subject 20217R is due to shadows cast by an overlying retinal blood vessel, which preclude reliable spectral classification by optoretinography. (**B**) Histograms showing the distribution of local proportion L selected for testing in each subject. The white-filled bars represent the distribution of test sites selected manually by the experimenter. The red lines and pink shaded areas show the results (mean ± 1 STD) of a Monte Carlo simulation (10,000 iterations) in which an equivalent number of test sites in each subject (n = 32 for 10001R; n = 26 for 20217R) was selected at random and histograms were recomputed following the same procedure.

### Retinal imaging and stimulation with adaptive optics

We measured the relationship between increment sensitivity and cone topography by stimulating retinal loci that differ in constituent L and M cone number. Deliberate stimulus placement at designated points of the photoreceptor mosaic required taking high resolution images of the retina, which we accomplished with an adaptive optics scanning laser ophthalmoscope (AOSLO). Our AOSLO, which is described in greater detail elsewhere (Mozaffari et al., 2020), raster scanned light from four channels onto the retina. These channels were: a 940 nm channel for wavefront sensing, an 840 nm channel for imaging, and two stimulation channels – 680 nm (± 14.9 nm) or “red” and 543 nm (± 13.5 nm) or “green”. The vergences of each channel were adjusted to counteract the longitudinal chromatic aberration of the typical human eye, as approximated by equation 5a of Atchison & Smith (2005). Two scanning mirrors, a 16 kHz horizontal resonant scanner and a 30 Hz vertical galvanometer scanner, produced the raster scan and hence set the 30 Hz frame rate of the display.

Photomultiplier tubes detected light returning from the retina, which was sampled at 20 MHz during the middle 80% of the forward scan to generate 512 x 512 pixel images. In the present experiment, the imaging field of view was about 0.9 degrees, and thus 1 pixel subtended ∼0.11 arcmin. This is about a quarter of the imaging channel’s diffraction-limited point spread function bandwidth, assuming a 6.5 mm pupil.

To draw stimuli at desired locations on the retina, acousto-optic modulators varied the intensity of the 543 nm or 680 nm light at appropriate points of the scan. Transverse chromatic aberration (TCA), which would normally cause stimuli of different wavelengths to be laterally offset from one another on the retina, was measured using an image-based method (Harmening et al., 2012), and corrected by modulating the 543 nm and 680 nm light at different times within laser scan. The subject used a bite bar during imaging which along with an X/Y/Z translation stage served to align the subject’s eye to the pupil plane of the system. To avoid changes in TCA with pupil position (Boehm et al., 2019), the experimenter used the translation stage to negate deviations of the pupil from its position at the time of TCA measurement, which was marked in a video feed of the subject’s 1^st^ Purkinje image. Prior to imaging, the subject’s eye was instilled with 1% tropicamide and 2.5% phenylephrine to dilate the pupil and freeze accommodation. A 1-second (= 30 frames) video of the retina was recorded for each trial of the experiment.

### Stimuli

All stimuli were squares, 2.25 arcmin in side length and 3 AOSLO frames (∼67 ms) in duration. Stimuli consisted of narrowband light centered on either 543 nm or 680 nm, and were shown against a large achromatic field, which was made of light from a Maxwellian-view RGB projector in addition to the infrared light used for imaging and wavefront-sensing. Light leak from the stimulation channels contributed negligibly to the background. The projector furnished a 1.1 deg square patch, which the subject aligned with the imaging raster to render the infrared light invisible, as well as a dimmer circular surround subtending 18 deg. The projector provided a small black spot to aid the subject’s fixation. The experimenter could move the fixation marker so that the desired retinal location was centered in the imaging field of view. The luminance of the square background was 1702 cd m^-2^, and the luminance of the surround was 851 cd m^-2^.

Both background and surround were approximately metameric to equal-energy white (MacLeod-Boynton coordinates: r = 0.6643, b = 0.0144; MacLeod & Boynton (1979)) and functioned to saturate rod activity and to approximately equalize activity in L, M, and S cones. Before the start of the experiment, the subject focused the projector display by adjusting the position of a Badal lens.

### Stimulus delivery

Retinal locations for stimulation were chosen at the beginning of each experiment, and stimuli were locked in place at these locations during presentation. Stimulus stabilization on the retina entailed several steps. First, an image of the retina captured over a single frame interval was designated to serve as the initial reference frame. All subsequent frames were registered to the reference frame by a fast strip-based cross-correlation algorithm (Arathorn et al., 2007; Yang et al., 2010). This yielded a live video of the photoreceptors in which displacements and distortions of the retinal image due to fixational eye movements were removed. Next, two seconds of stabilized video were recorded, and the frames of this video were averaged together to generate a high signal-to-noise ratio image which supplanted the initial reference frame. The new reference image was compared to a previously taken image of the retina in which L, M, and S cones were labeled, and based on this comparison the experimenter chose coordinates in the reference image at which to deliver stimuli. The same process for video stabilization produced real-time eye motion estimates that were used to track the points in the reference image selected for testing, so that stimuli could be reliably drawn at the desired loci. Stimulus delivery locations were marked in videos recorded during each trial with small digital crosses. Only trials in which both the stimulus x-position and y-position were ≤0.75 arcmin from the tracked x and y-positions for every frame of the delivery sequence contributed to threshold estimates. Out of 6,012 trials with subject 10001R, 5,266 (87.59%) were included, and out of 5,200 trials with subject 20217R, 4,348 (83.62%) were included. Of the included trials, the standard deviation of the x-position of the stimulus was 0.09 arcmin, and the standard deviation of the y-position was 0.10 arcmin, less than the diameter of a cone photoreceptor at our parafoveal test eccentricity. Thirty-one loci were targeted in subject 10001R’s retina, and 26 loci were targeted in subject 20217R’s retina. At one location for subject 10001R, thresholds were measured twice to confirm repeatability. Thus, 32 pairs of thresholds (for 543 and 680 nm stimuli) were collected from subject 10001R, and 26 from subject 20217R. The white boxes overlaid on the mosaic images in **Figure 1A** mark the locations targeted within each subject’s eye. We compared, by Monte Carlo method, the distribution of tested L/M ratios with the distribution expected from a completely random selection of test locations. For each subject, *n* retinal locations were randomly picked, where *n* is the number of unique 543 (or 680) nm threshold measurements made for the given subject. The L cone proportions at the random locations were determined (see *Computation of proportion L* below) and allocated to 8 equal-width bins over the range 0 to 1. This process was repeated 10,000 times in total, and the mean number of locations in each bin, as well as the bin count’s standard deviation across all iterations, were computed. The Monte Carlo results are graphed together with the observed distribution of proportion L in **Figure 1B**.

### Psychophysics

Subjects completed a self-paced seen/not seen task to determine thresholds for detecting small flashes of 543 or 680 nm light on a white background. On each trial, the subject pressed a key to initiate stimulus presentation, and indicated with an additional keyboard press whether the stimulus appeared red, green, or achromatic, or if it was not seen. In the case of poor stimulus delivery (which might be caused by a microsaccade, for instance), the experimenter instructed the subject to repeat the stimulus presentation and respond anew. Trial-to-trial stimulus intensities were generated from a Bayesian, adaptive staircase procedure, QUEST+ (Watson, 2017). In the typical measurement block, two discrete retinal locations were selected for testing; the local L/M ratios of the test locations were not disclosed to the subject. At each test location, sensitivity and color appearance were assessed with two test wavelengths: 543 nm and 680 nm. For each wavelength and location, two staircases ran in parallel, with one staircase starting at 25% of the maximum intensity and the other at 75%. Thus, each measurement block consisted of 8 independent staircases; these staircases were randomly interleaved so that the subject could not predict the intensity, wavelength, or retinal locus of stimulation based on the preceding trial. Except for a few early experimental sessions, each staircase contained 25 trials and, on 20% of trials the stimulus intensity suggested by the QUEST+ algorithm was ignored and stimulus intensity was instead drawn at random from a uniform distribution on the interval between 0 and the maximum intensity. Because threshold intensities were typically a very small fraction of the maximum power level, this was practically equivalent to interspersing high intensity stimuli throughout the staircase. These random, high intensity stimuli served to maintain the subject’s attention, as well as to assess suprathreshold color judgements. Generally, 100 trials per location were collected for each of the two stimulus wavelengths, per session. Three to six locations were tested every session, corresponding to a total of 600 to 1200 trials. Weibull functions were fit to the staircase data with the Palamedes toolbox, and threshold was taken as the stimulus intensity that corresponded to a 50% frequency of seeing.

### Computation of proportion L

The assortment of L and M cones hit by the stimulus is characterized in this paper as “proportion L”: the number of illuminated L cones out of the total number of illuminated L and M cones. A simple way to count the number of stimulated cones would be to tally the number of cone centers that fall within the nominal boundary of the stimulus on the retina. However, due to diffraction and any residual aberrations not corrected by the AOSLO, the boundary of the proximal stimulus is ambiguous. Additionally, a single cone admits light over an extended area and not just a single point. Our calculation of proportion L accounted for these nuances. We represented the cone mosaic as an array of two-dimensional Gaussians, which approximated the cone aperture function (MacLeod et al., 1992). Each Gaussian had a full width at half maximum of 0.59 arcmin and a volume of one. The spatial distribution of light on the retina during stimulation was computed by convolving an intensity-normalized distribution of light defined by the shape of the stimulus with a diffraction-limited point spread function computed for a 6.5-mm pupil and subsequently aberrated by 0.05 D of defocus to account for any residual aberrations (Harmening et al., 2014). The point spread function’s volume was normalized to unity, and its width was determined by the central wavelength of the stimulus (i.e., 543 nm or 680 nm). The retinal light distribution was multiplied with the array of L, M, or S cone aperture functions, and the integral of this product gave the number of cones of a particular spectral type that received the stimulus. Therefore, the number of stimulated L (or M, or S) cones is related to the amount of light accepted by the L (or M, or S) submosaic. The proportion L reported for a given threshold measurement is the average proportion L across trials. In cases where not all stimulated cones were classified, the range of possible L proportions (assuming all the unclassified cones were either L or M) is given in addition to the proportion L computed from only the classified cones.

### Modeling

Sensitivity to briefly-presented small spots might be expected to grow with the aggregate cone excitations produced by the light. Based on this idea, we computed theoretical 680/543 sensitivity ratios for every local L/M ratio encountered experimentally. Theoretical sensitivity ratios were derived from a simple additive model which is described below.

#### Cone photoreceptor sensitivities

To obtain individual cone spectral sensitivities, we estimated each subject’s lens, macular pigment, and photopigment optical densities. Lens density was computed from the CIE (2006) formulae, which account for age-related lens yellowing. Macular pigment density spectra were derived by scaling the normalized Bone et al. (1992) spectrum by an estimate of the peak macular pigment optical density (MPOD) within an interval containing the tested eccentricity. We estimated peak MPOD between eccentricities e1 and e2 from the following equation:

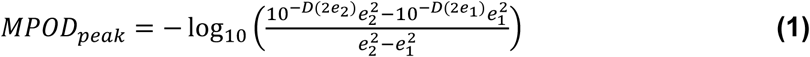

Where 𝐷(·) is the CIE (2006) function relating MPOD to field size. The radial distance between e1 and e2 was chosen to equal the diameter of the stimulus, with the tested eccentricity being the midpoint of the line connecting the two points. This approach presupposes radial symmetry in the distribution of macular pigment. The axial density of the photopigments in unbleached cones was calculated by multiplying the specific density provided by Bowmaker et al. (1978) by outer segment length measurements made by AO-OCT. The average outer segment length for subject 10001R was 35.7 μm, and that of subject 20217R was 32.9 μm. We estimated the amount of photopigment bleaching in the course of the experiment by computing the isomerization rates of L and M cones in response to the background light, and assuming a half-bleach constant of 10^6.4^ isomerizations s^-1^ (this is a conversion of the Rushton & Henry (1968) constant from trolands to isomerization rate for a standard 2:1 L/M ratio). This bleaching calculation was implemented through the *ComputePhotopigmentBleaching* routine in Psychtoolbox (Brainard, 1997).

#### Additive model

In our model, sensitivity is taken to be proportional to the response of a mechanism that combines L and M cone excitations. The responses to 680 and 543 nm stimuli at threshold are necessarily equal, and from this it can be shown that the ratio of 680 nm sensitivity (S_680_) to 543 nm sensitivity (S_543_) is given by the following expression:

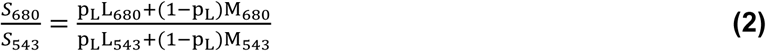

where p_L_ is proportion L, L_680_ (or M_680_) are the average L (or M) cone excitations per quantum of the 680 nm primary at the cornea, and L_543_ (or M_543_) are the average L (or M) cone excitations per quantum of the 543 nm primary at the cornea. Sensitivities are in terms of spectral power at the cornea (quanta s^-1^ nm^-1^), integrated over wavelength. L_680_, M_680_, L_543_, and M_543_ are computed from the custom cone sensitivities described in the preceding section.

#### Color appearance

As described above, the subject indicated whether the color appearance of each detected stimulus was best categorized as red, green, or achromatic. 680 nm light was seldomly categorized as “green”, and 543 nm light was rarely called “red” (see Results). Because of this, we opted to analyze the effect of stimulus intensity and cone topography on the probability of giving a color response consistent with the large-field appearance of the stimulus wavelength – that is, “green” for 543 nm and “red” for 680 nm. For convenience, we refer to these responses as “expected” color responses. In particular, we were interested in whether the subject was more likely to give the expected color response when the stimulus landed on spectrally-balanced regions of the cone mosaic, compared with regions dominated by a single cone class. For this analysis, we characterized the population of cones under the stimulus by its heterogeneity, defined as:

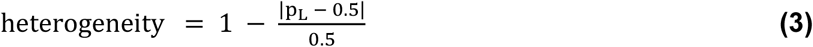

Where p_L_ is proportion L. This expression yields a heterogeneity value of 1 when the local L/M ratio is 1, and a value of 0 when the local receptor configuration is completely homogeneous (i.e. all L or all M).

We sought to examine the dependence of color naming on stimulus intensity, as well as two cone topographic features: L/M heterogeneity and distance to the nearest S cone (relative to the center of the targeted location). Intensity was expressed in terms of threshold multiple. The unit of S cone distance was cone spacing, computed for each subject by averaging the edge lengths of the Delaunay triangulation of cone locations. Using a simple linear regression was infeasible, because of the dichotomous response variable (on each trial, the subject either gave the expected color response or did not) and the non-independence of the observations (multiple trials per subject). Instead, we designed the following generalized linear mixed-effects model (GLMM):

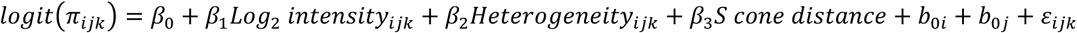

The probability of the expected color response from subject *i* at wavelength *k* is represented by 𝜋_𝑖𝑗𝑘_; 𝜀_𝑖𝑗𝑘_ is the residual term. Here, *i* is the subject index, *j* is the stimulus wavelength index, and *k* is the trial index for the *i^th^* subject and at the *k^th^* wavelength. Stimulus intensity, L/M heterogeneity, and S cone distance were treated as fixed effects with coefficients 𝛽_1_, 𝛽_2_, and 𝛽_3_ respectively. In addition to the fixed intercept 𝛽_0_, we accounted for between-subject variability by including the random intercept 𝑏_0𝑖_, and for differences in color responses between wavelengths with the random intercept 𝑏_0𝑗_. We fit the GLMM to our data with the Laplace method, using the *fitglme* routine (MATLAB 2024b; The MathWorks, Inc., Natick, MA, USA).

While a mixed-effects model best describes the structure of our data, our grouping variables (subject and wavelength) contained only two levels each, which may make it difficult to accurately estimate the random effects covariance parameters. We were satisfied that the inclusion of random effects was appropriate after taking the following steps. First, we confirmed that if subject and wavelength were treated as fixed (rather than random) effects, the coefficient estimates for intensity, heterogeneity, and S cone distance remained within 1% error of the corresponding estimates from the mixed effects model. Second, we confirmed by likelihood ratio test that the mixed-effects model outperformed a model which contained only intensity, heterogeneity, and S cone distance as fixed effects (λLR = 931.56, Δdf = 2, ΔAIC = -927.36, p < 0.001).

## RESULTS

### 680/543 nm sensitivity vs. local L/M ratio

Thresholds were measured for stimuli targeted to the retinal regions indicated in **Figure 1A**. Each flash stimulated an average of 4.6 cones in subject 10001R’s eye, and 4.0 cones in subject 20217R’s eye, with cone number calculated as described in the *Computation of proportion L* section. The histograms in **Figure 1B** summarize the local L/M ratios sampled throughout the study, in addition to the variety in L/M ratio that would arise from arbitrary stimulus placement. The empirical distributions of local proportion L we obtained for each subject were qualitatively similar to the simulated distributions that resulted from randomly selecting test locations. In other words, although we manually selected locations for testing, systematic sampling biases are unlikely to underlie the results we describe below. That being said, we did make a deliberate effort to increase our sampling of majority-M cone regions in subject 20217R, who has a higher-than-average global L/M ratio, and in the end targeted three such locations. According to the Monte Carlo simulation, the probability of encountering three or more majority M cone regions in subject 20217R’s eye, given the number of locations tested, was only ∼23%.

The spread of sensitivities to 543 nm and 680 nm flashes is depicted in the box plots of **Figure 2A**. The solid lines show luminosity function predictions for each subject generated by weighting the subject-specific cone fundamentals according to the relative density of L and M cones in the classified mosaics (see **Figure 1A**). The difference between the median sensitivities to 543 and 680 nm small spots was 1.68 log10 units for subject 10001R, and 1.80 log10 units for subject 20217R, which is in qualitative agreement with the V(λ) functions. The interquartile range (IQR) of 543 nm sensitivities spanned 0.12 log10 units for subject 10001R and 0.16 log10 units for subject 20217R, while the IQR of 680 nm sensitivities were slightly broader, spanning 0.22 log10 units for subject 10001R, and 0.34 log10 units for subject 20217R. Since L and M cones have almost identical sensitivities to 543 nm light, the dispersion of 543 nm thresholds must have been driven primarily by factors other than cone topography. In contrast, L cones absorb more at 680 nm than M cones, and so the increased variability in our observers’ 680 nm thresholds likely originated, at least in part, from local fluctuations in L/M ratio. In addition to this within-observer variability, we note that, for both stimulus wavelengths, subject 10001R was more sensitive than subject 20217R, likely because the latter was tested at a slightly more eccentric location where cone density is lower (**Figure 1A**). Thus, to examine the impact of proportion L on sensitivity while discounting other factors, we normalized the 680 nm sensitivity measured at each test locus by the corresponding 543 nm sensitivity.

**Figure 2.**
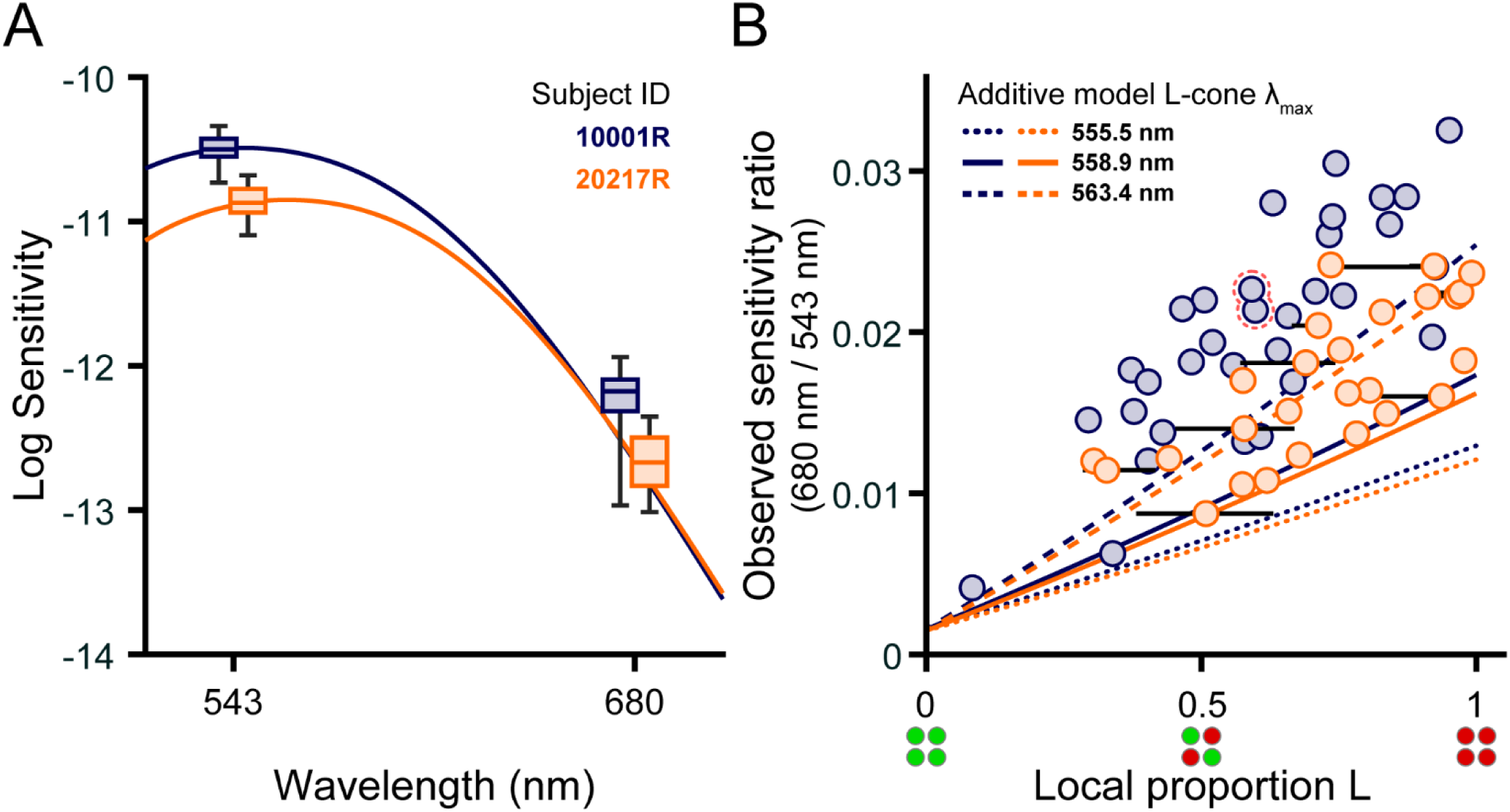
Local cone topography influences small-spot spectral sensitivity. (**A**) Boxplots of sensitivity to 543 nm and 680 nm narrowband increments obtained from subjects 10001R (blue) and 20217R (orange). Boxes indicate the median sensitivity (middle line) and interquartile range (upper and lower box edges); the whiskers span the full range of sensitivities measured with each wavelength. For clarity, each subject’s boxes have been shifted ±5 nm from the tested wavelength. Retinal locations selected for testing are shown in Figure 1. Solid lines indicate predicted luminous efficiency functions, V(λ), computed for each subject as described in the text; the V(λ) functions are laterally shifted ±5 nm and vertically positioned to best fit the empirical data. Sensitivities are defined as the reciprocal of the detection threshold intensity expressed in units of quanta s^-1^ deg^-2^ at the cornea. (**B**) The ratio of 680 nm sensitivity to 543 nm sensitivity is plotted against the local proportion L of the stimulated region (see Methods). The blue and orange circles correspond to data from subjects 10001R and 20217R, respectively. In instances where the stimulated region contained an unclassified cone, the range of possible local L/M ratios (calculated assuming the missing cone is either L or M) is denoted by a black horizontal line. The pair of points with a dashed red outline correspond to the locus selected for repeat testing in subject 10001R. Predictions from the additive model described in the text are shown by the blue (10001R) and orange (20217R) lines, where the line format indicates the λmax value of the L-cone pigment used to model sensitivity (Kraft et al., 1998).

The ratio of 680 to 543 nm sensitivity is plotted as a function of local proportion L in **Figure 2B**. Subject 10001R’s data are represented by the blue circles, and subject 20217R’s by the orange circles. As expected, normalized sensitivity to 680 nm increased with relative L cone numerosity, and significant correlations between these variables were found for both subjects (10001R: Spearman’s rho = 0.80, p < 0.001; 20217R: Spearman’s rho = 0.73, p < 0.001). There was considerable variability in thresholds measured for patches of retina with similar proportions of L cones; this variability would preclude precise estimation of L/M ratio from detection thresholds. Data from subject 10001R generally lie above data from subject 20217R, indicating that the former subject is more sensitive to 680 nm stimuli than the latter, all else being equal.

This intersubject variability may reflect differences in L cone photopigments. The dotted, solid, and dashed curves give the 680/543 sensitivity functions predicted from **Equation 2**, assuming L cone spectral peaks at 563.4, 558.9, or 555.5 nm. The predictions for subjects 10001R (blue) and 20217R (orange) differ slightly due to interobserver differences in cone outer segment length (see ***Cone photoreceptor sensitivities*** section above). Data from subject 10001R mostly fall above the additive model predictions, while data from subject 20217R fall near the curves for which the L cone λmax is either 558.9 or 563.4 nm. This does not necessarily mean that subject 10001R’s L cone spectral peak lies further in the red than would be explained by known L cone pigment variants. Instead, it may be that the additive model underestimates long-wavelength sensitivity by failing to account for signals from non-additive (i.e., cone-opponent) mechanisms that may contribute to detection.

### Comparison of predicted and observed data

Figure 3A plots, for each targeted location, the 680/543 nm sensitivity ratios predicted by the additive model against the observed ratio. In this plot, predictions for subject 10001R (blue markers) were made assuming an L cone λmax of 563.4 nm, while a λmax of 558.9 nm was used for subject 20217R (orange markers). The use of different spectral peaks was meant to account for the systematic differences between the two subjects’ data (Figure 2B), though it is entirely possible that both subjects have the same L cone opsins, and the enhanced long-wavelength sensitivity of subject 10001R might be explained by post-receptoral factors. Most of the data in Figure 3A lie close to the 1:1 line, suggesting that our empirical results are consistent with the hypothesis that small spot detection is largely mediated by a mechanism that adds unweighted inputs from L and M cones in its receptive field; however, the theoretical data tend to underestimate the observed sensitivity ratios.

**Figure 3.**
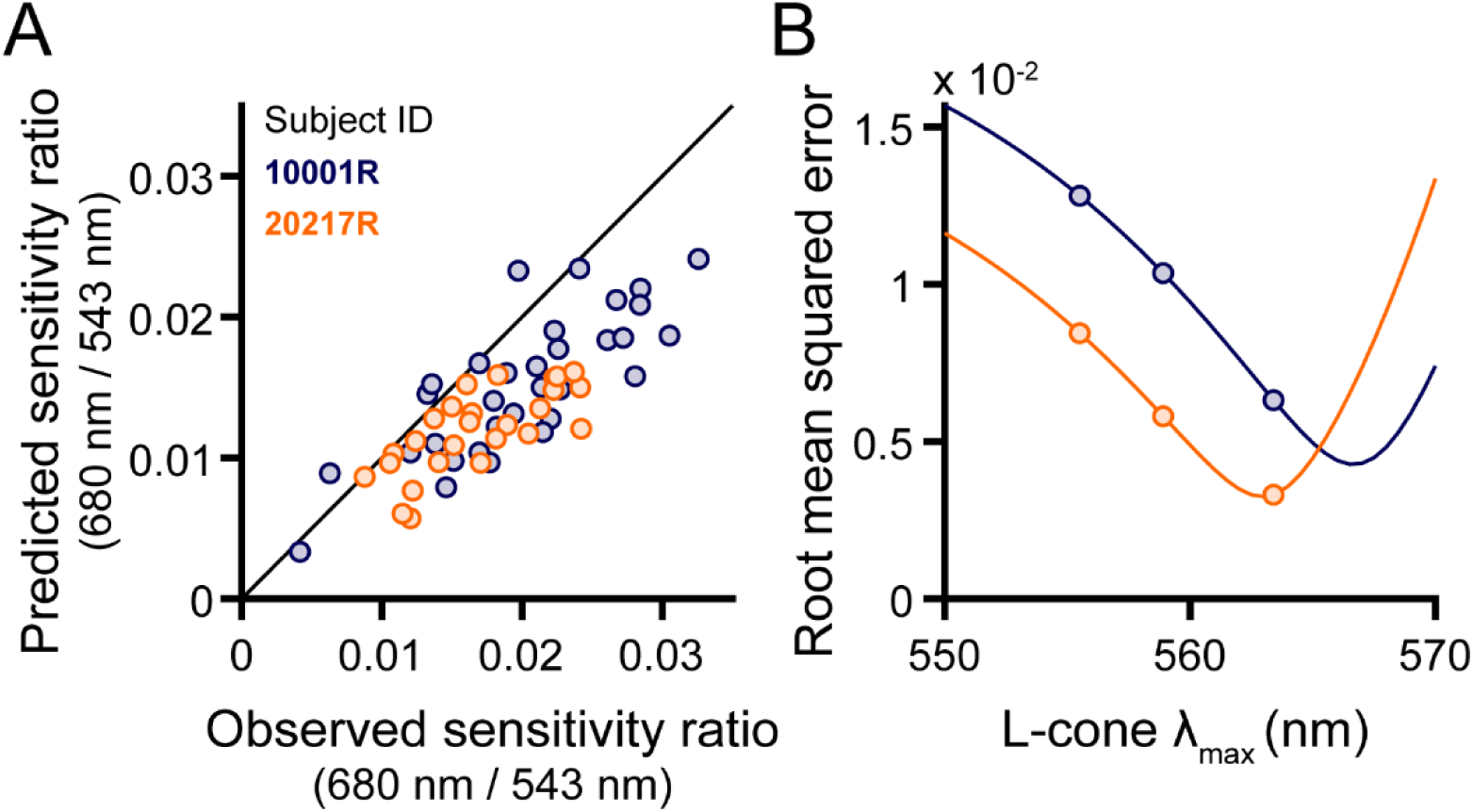
Modeling the 680/543 nm sensitivity ratio. (**A**) Predicted 680/543 nm sensitivity ratios are plotted against their empirically-measured counterparts. Additive model predictions for each retinal locus are indicated by circles. For subject 10001R, an L-cone λmax of 563.4 nm was used to generate the predicted data; for subject 20217R, a value of 558.9 nm was chosen. (**B**) Root mean squared errors (RMSE) of the additive model predictions are plotted as a function of L-cone λmax for subject 10001R (blue) and 20217R (orange). Markers are shown at three physiologically-plausible values of L-cone λmax for both models. In addition, solid lines depict the performance of the additive model evaluated with finer L-cone λmax sampling, revealing minima at or beyond the longest values reported in the literature.

To understand how errors in the model predictions vary with the choice of L cone λmax, root mean square error (RMSE) is graphed as a function of λmax in Figure 3B. In this plot, the continuous lines give the additive model RMSE over a range of wavelengths. The errors for the additive model at three biologically plausible spectral peaks are plotted as circles. If selection is restricted to the three λmax values of interest, the RMSE is similar between the two subjects when a wavelength of 563.4 nm is selected for 10001R, and a wavelength of 558.9 nm for 20217R. However, as the solid curves indicate, the RMSE approach minima at longer wavelengths than those just stated: the minimum for 10001R occurs around 566.5 nm, and the minimum for 20217R occurs around 562 nm. Thus, to best model the data for subject 10001R, the L cone sensitivity peak chosen must exceed the maximum value reported in the literature (∼563 nm). The difference between the optimal additive model λmax values for the two subjects is ∼4.5 nm, which is similar to the difference between the two most commonly found L-cone spectral peaks (Kraft et al., 1998; Neitz & Neitz, 2011).

### Color appearance

Our color appearance results are summarized in Figure 4. The stacked bar plots in Figure 4A represent the proportion of seen trials categorized as red, green or achromatic. For both stimulus wavelengths, detected flashes were most frequently categorized as achromatic (90.5% for 543 nm; 58.2% for 680 nm). The long-wavelength stimulus was perceived as red at a higher rate (47.7% for 10001R; 32.5% for 20217R) than the middle-wavelength stimulus was as green (6.6% for 10001R; 9.1% for 20217R). “Non-expected” color responses were rare: 680 nm flashes were almost never seen as green (0.06% for 10001R; 1.01% for 20217R), and the 543 nm stimuli were very seldom called red (0.44% for 10001R; 4.01% for 20217R). Of note, subject 20217R, whose retina is comparatively L-rich, was less likely to call 680 nm stimuli red.

**Figure 4.**
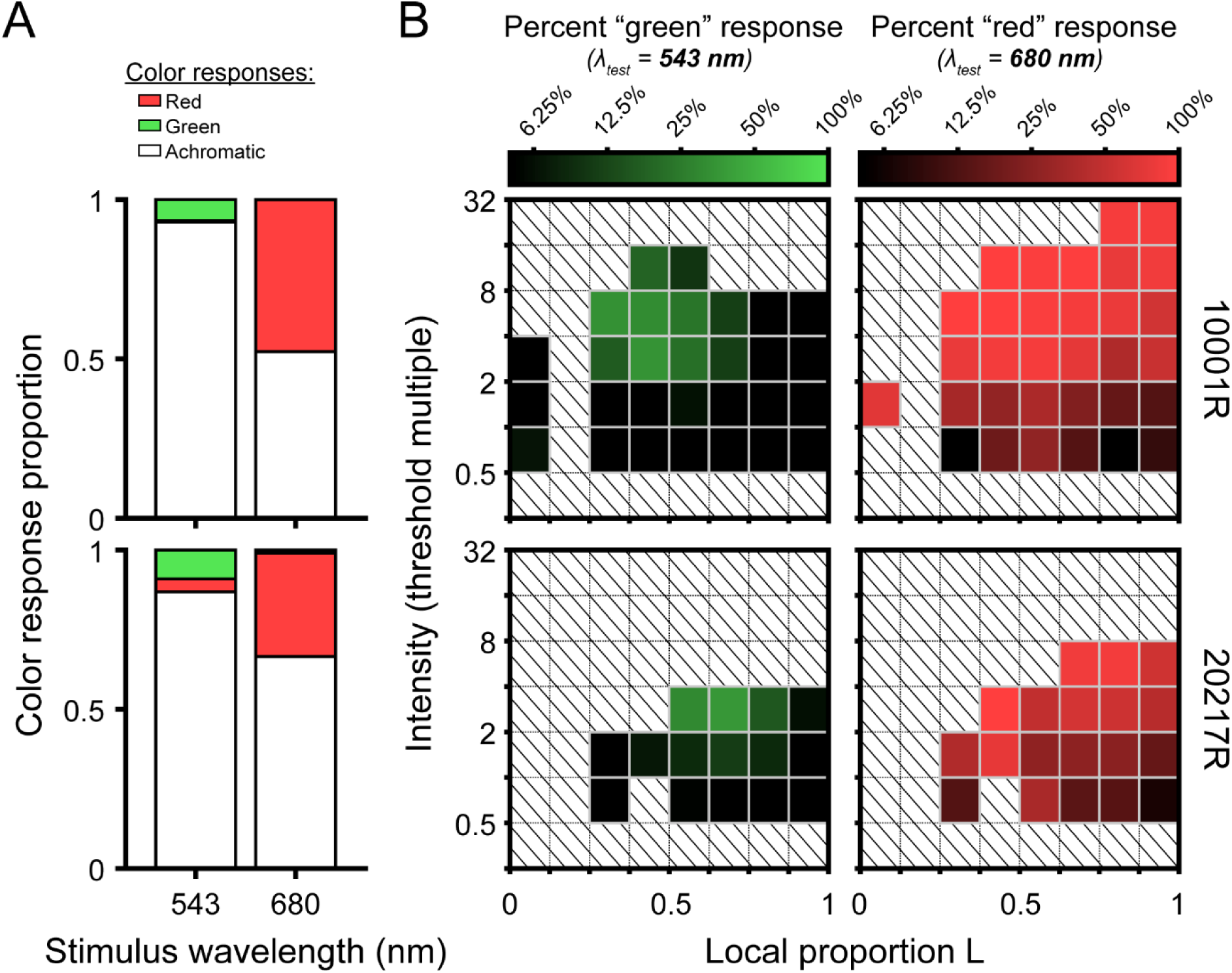
Summary of small-spot color appearance results. (**A**) Stacked bar plots showing the proportion of seen trials categorized as red, green, and achromatic as a function of stimulus wavelength. Results from subjects 100001R and 20217R are shown in the top and bottom rows, respectively. (**B**) Color-coded matrices represent the naming data from (A) binned by local mosaic demographics and stimulus intensity (relative to local threshold). Darker squares indicate intensity-mosaic combinations that were more likely to yield achromatic percepts, whereas brighter green and red squares indicate a higher likelihood of responding “green” and “red” to the 543 nm and 680 nm flashes, respectively. Note that “non-expected” color responses (e.g., categorizing a 680 nm stimulus as “green”) occurred rarely and were thus excluded from these plots. To enhance visualization, bins containing fewer than 10 trials are not shown. Squares filled with diagonal lines denote empty bins.

To examine the effects of local cone topography and stimulus intensity on color appearance, the data were binned along the intensity and local proportion L dimensions, and for each bin, the fraction of trials on which the subject gave the expected color response was calculated. The matrices in Figure 4B give the frequency of expected color responses as a function of intensity and L cone proportion. Darker matrix elements correspond to high rates of achromatic percepts. In these plots, intensities are expressed as multiples of the local threshold, which has implications for the distribution of matrix bins. As described in the Methods, trial intensity was primarily determined by an adaptive staircase, which concentrates most presentations near threshold; however, on a random subset of trials, high-intensity stimuli were interspersed.

Because subject 20217R had lower absolute sensitivity to both stimulus wavelengths (Figure 2A), these high-intensity trials end up being allocated to intensity bins closer to threshold. In addition, 680 nm sensitivity increased with local proportion L in both subjects (Figure 2B).

Consequently, 680 nm flashes of a fixed intensity would be assigned to intensity bins increasingly farther from threshold as the local proportion L increases. Despite this asymmetry in stimulus detectability across observers, wavelength, and retinal locus, Figure 4B suggests the likelihood of calling 680 nm flashes red or 543 nm flashes green grew with normalized stimulus intensity.

The effect of local L/M ratio on color naming was less obvious than that of intensity. For the 543 nm stimulus, green responses were almost never given for stimuli falling within the leftmost or rightmost bins, which correspond to highly asymmetric L/M ratios (Figure 4B, left column).

Instead, green responses were concentrated near 50% L at higher stimulus intensities. If a similar hotspot exists for 680 nm, it is more subtle. At near-threshold intensities, chromatic responses appear to be more frequent for retinal loci with balanced L and M numbers, but the color response matrix rows become more uniform as normalized stimulus intensity increases (Figure 4B, right column). In subject 10001R, one retinal locus containing mainly M cones was tested (Figure 4B, top row, leftmost matrix columns). With 543 nm light, predominantly achromatic percepts were logged at this spectrally homogenous location across a range of stimulus intensities, mirroring the pattern observed in more frequently encountered regions dominated by L cones. For 680 nm light, however, the stimulus was reliably categorized as “red”. We suspect this may be a consequence of the higher absolute illuminance required to detect 680 nm flashes landing in this M-rich region, leading to increased light scatter onto nearby cones that might counteract the decline in chromatic sensitivity that would otherwise occur as local proportion L approaches zero.

To test the hypothesis that L/M ratio symmetry would be advantageous for color vision (Gunther & Dobkins, 2002; Hood et al., 2006), we categorized each stimulated retinal locus by its spectral heterogeneity, applying an index similar to the one developed by Hofer, Singer, et al. (2005) to quantify the symmetry of global L/M cone ratios (see **Equation 3**). A schematic illustrating the relationship between local spectral composition and the heterogeneity index is shown in Figure 5A. The distribution of heterogeneity indices encountered across all trials is depicted in Figure 5B. We divided this distribution into tertiles to visualize the joint effects of mosaic heterogeneity and normalized stimulus intensity on the probability that the subject gives a chromatic response. In Figure 5C, it is clear that increasing stimulus intensity (relative to threshold) increases the likelihood of categorizing the stimulus as chromatic, reiterating the trends we observed in the color response matrices of Figure 4. Comparing the intensity-response curves for the three heterogeneity bins, it is also apparent that the likelihood of our small test flash being judged as chromatic increases as the mosaic becomes more heterogeneous – at least for stimulus intensities at and slightly above threshold.

**Figure 5.**
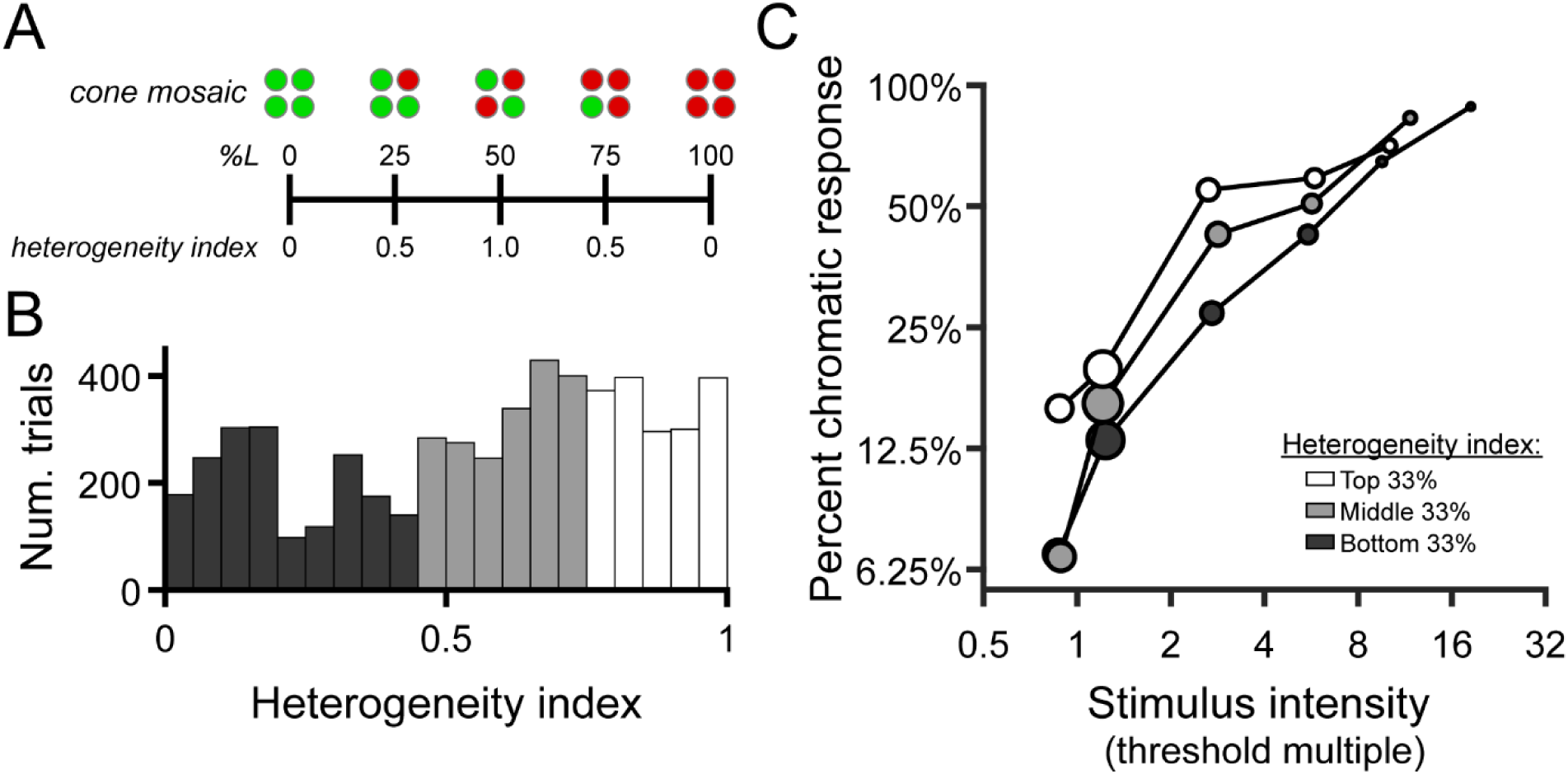
Chromatic responses increase with stimulus intensity and mosaic heterogeneity. (**A**) Schematic depicting the relationship between local cone configurations and the heterogeneity index described in the methods. (**B**) Histograms showing the distribution of heterogeneity indices encountered across all seen trials (n = 5,552). Bars are color-coded to represent the bottom (dark gray), middle (light gray), and top (white) tertiles of the heterogeneity distribution. Data from the two study participants are combined in this analysis. (**C**) The percentage of trials in which the “expected” chromatic response (“red” to 680 nm or “green” to 543 nm) was given is plotted as a function of binned stimulus intensity and mosaic heterogeneity. As in Figure 4, stimulus intensity is expressed in multiples of local detection threshold; the tick marks on the abscissa indicate the edges of the intensity bins. Markers are color-coded according to the heterogeneity tertiles shown in (B). Marker area is scaled to the number of trials in each bin, and its x-position represents the median stimulus intensity within each bin.

We determined whether changes in intensity, L/M heterogeneity, and S cone distance significantly changed the odds of responding “red” to 680 nm stimuli or “green” to 543 nm stimuli by fitting a GLMM to our data (as described in Methods). The effects of log2 intensity (β1) and L/M heterogeneity (β2) were significant (both with p < 0.001). Doubling the stimulus intensity was predicted to increase the odds of giving the expected color response by a factor of 2.94 (95% CI: [2.72, 3.18]). A unit increase in L/M heterogeneity (i.e., a change in proportion L from 0 or 1 to 0.5) was predicted to increase the same odds by a factor of 5.51 (95% CI: [4.11, 7.40]). The model predicted no effect of S cone proximity on the odds of giving a chromatic response (odds factor: 1.01; 95% CI: [0.92, 1.10]; p = 0.889). The adjusted R^2^ for the GLMM was 0.4458, which suggests that a little under half of the variation in the empirical data is explained by the model. The full GLMM outperformed a reduced model in which the heterogeneity term was removed (λLR = 139.31, Δdf = 1, ΔAIC = -134.31, p < 0.001), further supporting the importance of cone ratio in small-spot color perception.

## DISCUSSION

Small spot detection, discrimination, and color naming data have been used before to make inferences about the distribution of L, M, and S cones in the retina as well as the nature of mechanisms that combine inputs from cones of different type (Chaparro et al., 1994; Cicerone & Nerger, 1989; Finkelstein & Hood, 1984; King-Smith & Carden, 1976; Krauskopf, 1964; Otake et al., 2000; Vimal et al., 1989). Distinguishing the influence of cone spectral demographics from the effects of postreceptoral processing on behavioral data is, however, challenging in the absence of objective measurements of cone numbers (e.g., Krauskopf, 2000). We studied the direct link between cone mosaic organization and behavior by measuring small spot detectability and color appearance in subjects whose cones were spectrally classified by an objective method. Our stimuli were delivered through an AOSLO, which mitigated the effects of optical aberrations and fixational eye movements, in turn enabling us to estimate the trichromatic inputs to post-receptoral mechanisms on each experiment trial. Our results permit two conclusions to be drawn about the relationship between local cone mosaic topography and vision. First, as expected, spectral sensitivity to short-duration, retinally-stabilized small spots depended strongly on local L/M cone ratio. Second, subjects were more likely to rate brief, monochromatic flashes as chromatic if they were sampled by a balanced number of L and M cones. We discuss these findings in turn below.

### Local L/M cone ratio and small-spot detection

The dependence of visual sensitivity on cone spectral topography was relatively straightforward: long-wavelength stimuli were overall more easily seen as the retinal patches they landed on became increasingly L cone rich (Figure 2B). The relationship between the 680/543 nm sensitivity ratio and local L/M ratio was reasonably well-described by a non-opponent detection model that included a characterization of early visual processing through the photoisomerization stage (Figure 3A). Together, these results are compatible with the idea that carefully collected behavioral data can reveal spatiochromatic variations in the receptor array (Cicerone & Nerger, 1989; Krauskopf, 1964; Otake et al., 2000; Vimal et al., 1989; Williams et al., 1981). However, despite the strong correlation between our measurements of local spectral sensitivity and objectively-determined cone topography, several factors evident in our data would complicate the precise determination of local L/M ratio solely from detection thresholds as measured in this study.

First, as can been seen in Figure 2B, locations within an individual retina with similar L/M numerosity are not always equally sensitive. The origins of this within-subject variability are not entirely clear. It likely reflects a combination of neural and psychophysical factors. Regarding the former, physiological evidence from the primate retina suggests that downstream neurons may not sample L and M cones with uniform weights across small spatial scales, even within retinal ganglion cell subclasses that do not systematically favor one cone type over the other (Field et al., 2010; Li et al., 2014). Likewise, it seems plausible that perceptual judgments reliant on the signals of fewer than a half-dozen cones will be inherently noisy. Although the one test site we deliberately retested in separate sessions yielded near-identical sensitivity ratios (Figure 2B, outlined pair), understanding how much variance in our data might be attributed to psychophysical noise would require a more comprehensive analysis of test-retest variability in our experimental paradigm. Finally, an additional source of experimental noise could arise from imperfect cone spectral classifications. In our measurements, most of the stimulus light fell on fewer than five cones on any given trial, and so a single misclassified (or unclassified) cone in the stimulated region would produce significant error in our objective estimate of the underlying L/M ratio. Given that previous work has estimated the error rate of OCT-based cone classification methods to be less than 0.5% (Pandiyan et al., 2022; Zhang et al., 2019), we expect spectral misclassifications to contribute only negligibly to the variance present in our dataset.

Intersubject variability poses a second, arguably more substantial challenge to estimating proportion L from increment sensitivity measurements. In our sample of two trichromats, one subject (10001R) tended to exhibit higher 680/543 nm sensitivity ratios than the other (20217R) when the tested locations had matched L/M ratios (Figure 2B). This aligns with previous work articulating how factors other than cone numerosity can drive considerable between-subject differences in spectral sensitivity (Bieber et al., 1998; He et al., 2021). The greater sensitivity ratio of subject 10001R is partially explained by longer cone outer segments at the eccentricity examined for this subject, compared with subject 20217R. Aside from cone outer segment length, a well-known source of individual differences in the color-normal population originates from genetic variations in the L opsin gene that alter its spectral tuning (Kraft et al., 1998; Neitz & Jacobs, 1986; Neitz & Neitz, 2011). In the non-opponent modeling of our observers’ data, comparable amounts of variance in the data could be accounted for by shifting 10001R’s L-cone λmax ∼4.5 nm beyond the value used for 20217R (Figure 3B). In addition to differences in cone opsin genetics, trichromatic encoding can be affected by pre-retinal filtering by the lens and macular pigment (Asano et al., 2016). While genetic sequencing can be used to estimate the peak sensitivities of the L and M cone opsins (Neitz & Neitz, 2011), psychophysical approaches based on color matching have been proposed to derive more comprehensive estimates of cone action spectra in individual eyes (Shi et al., 2024; Thomas & Mollon, 2004). Applying these techniques to build more realistic models of our subjects’ early visual systems is a goal of future work.

The additive model (**Equation 2**) assigns equal weight to individual L and M cones in the mosaic. Hofer, Carroll, et al. (2005) compared L/M ratios estimated by retinal densitometry and flicker electroretinography (ERG) in a group of subjects whose opsin genetics had been characterized. They observed a discrepancy between the two methods that could be reconciled if the M-cone contribution to the mechanism(s) engaged by flicker ERG was ∼1.5-folder higher than that of L cones. Adjusting the L and M cone weights in our model accordingly did not produce better fits, with the model RSMEs at the three physiologically plausible values of L-cone λmax increasing by an average of 13.4 ± 7.3% (mean ± STD; one-sided Wilcoxon signed rank test: T = 21, p = 0.0156, N = 6). Although it remains possible there could be subtle cone-type-specific differences in L/M input to non-opponent pathways, it would be difficult to disambiguate such effects from the other sources of variability described above.

Finally, we note that, alongside pre-retinal and receptoral factors, the involvement of non-additive post-excitation mechanisms could further obscure the link between cone numbers and small-spot detection thresholds. Although the systematic increase in sensitivity to long-wavelength light with local L cone density comports with a significant contribution from non-opponent mechanisms to small-spot detection, our additive model predictions tended to underestimate relative sensitivity to 680 nm across a range of L cone λmax values examined (Figure 2B; Figure 3A). Similarly, although the majority of near-threshold stimuli generated achromatic percepts (Figure 4A), slightly suprathreshold stimuli were often categorized as “chromatic”, particularly when the stimulus wavelength was 680 nm (Figure 4B). These facts collectively point towards a secondary contribution from an opponent channel to our sensitivity measurements.

Although studies using extended grating stimuli generally find the spatiotemporal resolution of chromatic mechanisms to be lower than their achromatic counterparts (Lee et al., 1990; Mullen, 1985; Sekiguchi et al., 1993; Wuerger et al., 2020), the idea that small, brief flashes can engage both pathways is supported by detection and discrimination data collected with small-field increments. King-Smith & Carden (1976) found evidence that spectral sensitivity to small (3 arcmin), brief (10 ms) increment flashes presented on an intense neutral background resembles the photopic luminous efficiency function measured with larger stimuli via heterochromatic flicker photometry. This concordance suggests the involvement of a non-opponent, “achromatic” pathway in detecting narrowband increments of small angular subtense. However, their results also indicated that, at longer wavelengths, near-threshold stimuli could be readily discriminated from white, leading them to conclude that the so-called “color” and “luminance” systems have similar sensitivities in this part of the spectrum. By contrast, King-Smith and Carden’s results indicated a substantial reduction in the sensitivity of the chromatic system to minute flashes of intermediate wavelength, presumably due to the similarity in L and M responses produced by mid-spectrum lights. Subsequent investigations into small-spot detection and discrimination also support a role for “red-green” opponent mechanisms in processing small flashes near threshold, particularly for longer wavelength stimuli (Chaparro et al., 1994; Finkelstein & Hood, 1984).

The likely involvement of the L-M pathway in our detection data introduces another plausible explanation for the sensitivity differences we observed between individuals. Suprathreshold color appearance has been shown to be robust to large individual variations in L/M ratio (Brainard et al., 2000; Miyahara et al., 1998; Neitz et al., 2002). One way to accomplish this invariance would be to reduce the input gain of the more numerous cone type to downstream cone-opponent circuits. In cases where L cones predominate, such sensory regulation would reduce the relative contribution of the chromatic pathway to detection. The loss of chromatic sensitivity could manifest as a general elevation in long-wavelength thresholds and a corresponding increase in “achromatic” percepts near threshold. In line with this hypothesis, we found the subject with a greater abundance of L cones (20217R; L/M ratio = 3.66) exhibited a lower relative sensitivity to long-wavelength stimuli (Figure 2) and was less likely to categorize 680 nm flashes as “red” (Figure 4) in comparison to subject 10001R, whose L/M ratio of 1.47 is closer to the typical value (Carroll et al., 2002). While the mechanistic basis of these trends remains unclear, obtaining a more detailed characterization of the light encoding properties of our subjects’ cone mosaics would, as described above, enable the relative contributions of post-receptoral processes to our measurements to be delineated more clearly.

### Cone mosaic topography and small-spot color appearance

#### Local L/M cone ratio

In addition to measuring detection performance, our observers were asked to assert whether the appearance of each seen trial was best described as “red”, “green”, or “achromatic”. While this tripartite categorization scheme may seem coarse in relation to the richness of human color experience, previous color naming studies suggest these terms capture the majority of sensations evoked by small middle- and long-wavelength flashes presented on a neutral photopic background (Bouman & Walraven, 1957; Cicerone & Nerger, 1989; Krauskopf & Srebro, 1965; Sabesan et al., 2016; Schmidt, Boehm, et al., 2018). As shown in Figure 4A, we found a disparity in the chromatic response rates for the 543 nm and 680 nm flashes, with the latter stimuli more frequently rated as “red” than the former were categorized as “green”.

These differences likely arise from the relative insensitivity of the L-M pathway to middle-spectrum lights of small angular subtense, as discussed in the preceding section (Chaparro et al., 1994; King-Smith & Carden, 1976). Despite the asymmetry in overall response rates to our two narrowband stimuli, our analysis demonstrates that flashes of either wavelength were more likely to be categorized as “chromatic” when the retinal neighborhood they encountered had an L/M cone ratio near unity (Figure 5). Because the mosaic dependence of color naming was maintained when stimuli were equated for detectability, we interpret our appearance results as evidence that the sensitivity of the “red-green” chromatic mechanism covaries with the underlying distribution of L and M cones.

Prior studies have produced data in support of the idea that L/M ratio symmetry may be beneficial for chromatic detection and appearance mechanisms. Gunther & Dobkins (2002) found a relationship between L/M ratio estimated from heterochromatic flicker photometry and sensitivity to red/green gratings, concluding that observers with ratios closer to one had superior chromatic contrast sensitivity compared to observers with skewed ratios. In subjects whose mosaics had been characterized using adaptive optics densitometry, Neitz et al. (2020) also observed a positive correlation between L/M ratio symmetry and sensitivity to fine-grained red/green gratings, mirroring earlier results suggesting the ability to localize the gap in cone-isolating “Landolt C” targets may depend on the relative numbers of L and M cones (Danilova et al., 2013). Finally, Hofer, Singer, et al. (2005) reported that individuals with more balanced L/M ratios were more likely to rate cone-sized flashes as chromatic. Our results demonstrating a direct relationship between local cone demographics and color appearance are in keeping with these earlier findings, although it is worth noting that our study has the added advantage of knowing both the spectral topography of the observer’s cone mosaic and the landing position of the stimulus on each trial.

A spectrally heterogeneous mosaic may not only be beneficial for chromatic sensitivity, but also for discriminating color from luminance information. In the context of the present experiment, the task of distinguishing a small, monochromatic light from an equally luminous broadband (white) light must be accomplished using signals from a small number of cones. If the stimulated receptor array was composed of a single spectral type, identical sets of cone excitations would be generated by the monochromatic and white lights, and the two stimuli should, in principle, be indistinguishable.^1^ If one constituent in the local receptor neighborhood were replaced with a cone of another type, then monochromatic and white lights could be discriminated on the basis of the outlier cell’s response, though presumably noise in the underrepresented spectral class would limit performance (Vorobyev & Osorio, 1998).

Measurements made from carriers of X-linked color vision deficiencies indicate that red-green chromatic discrimination is impaired individuals with skewed cone ratios (Hood et al., 2006). In a similar vein, although the midpoint of the classical Rayleigh match should depend solely on the spectral sensitivity of the L and M cones, it has been hypothesized that individuals with asymmetric L/M ratios will accept matches over a broader range, presumably because their discriminative capacity is diminished (Barbur et al., 2008; Zhaoping & Carroll, 2016). While we did not formally measure chromatic discrimination thresholds in the present study, it is evident from comparing the color response matrices in Figure 4B that our 543 nm and 680 nm stimuli could readily be distinguished at most intensities and retinal loci. Elucidating whether the apparent advantage of L/M symmetry for chromatic sensitivity also extends to chromatic discrimination is a goal of ongoing work.

An alternative approach to identifying what, if any, L/M cone ratio is optimal for color vision is to evaluate the degree to which the signals encoded by the cone mosaic can be used to infer the properties of the distal stimulus. In addition to incorporating contributions from receptor noise, such reconstruction frameworks enable the role of L/M numerosity and arrangement to be evaluated in the presence of other factors thought to be important for perceptual inference, including natural image statistics, physiological optics, and prior visual experience. Using a Bayesian approach to reconstruct signals from a simple dichromatic retina, Manning & Brainard (2009) concluded that a spectrally balanced cone mosaic produces optimal estimates of the retinal image, provided the noise characteristics of the two receptor classes are similar.

Subsequent computational investigations into mosaic optimality found that the advantages conferred by a symmetrical L/M ratio become less pronounced once longitudinal chromatic aberrations are incorporated into the simulation (Garrigan et al., 2010; Zhang et al., 2022). The empirical findings we report here may serve as a useful benchmark for future attempts to identify the conditions under which the trichromatic cone mosaic does (or does not) constrain human color perception.

Whether the mosaic-dependent variation in chromatic sensitivity we observed at small spatial scales has implications for suprathreshold color perception remains uncertain. Prior studies using large-field stimuli have generally found that certain aspects of color appearance, including the spectral locus of unique yellow, are invariant to individual differences in L/M ratio (Brainard et al., 2000; Jordan & Mollon, 1997; Miyahara et al., 1998; Neitz et al., 2002). This constancy could be achieved if downstream cone-opponent neurons amplify diminished L-M signals caused by asymmetric cone ratios. It has been shown that unique yellow can be manipulated by changing the spectral characteristics of the external environment (Neitz et al., 2002; Welbourne et al., 2015), lending credence to the notion that color-mechanisms calibrate themselves according to the statistics of their input signals (Mollon, 2006). Likewise, considerable attention has been paid in recent years to how such sensory regulation might account for the near-normal suprathreshold color vision of individuals with anomalous trichromacy (Boehm et al., 2014, 2021; Emery et al., 2023; Knoblauch et al., 2020; Tregillus et al., 2021). If similar processes are engaged by local variations in the spectral composition of the cone mosaic, one might expect the topography of color appearance to become more uniform as stimuli become increasingly suprathreshold.

#### Proximity to S cones

Although S cones would not be directly activated by the 543 and 680 nm test wavelengths we used in this study (Stockman et al., 1999), the presence of short wavelength receptors in the stimulated neighborhood could theoretically influence how the visual system interprets signals from nearby L and M cones. There are two possible explanations for how this influence may arise. First, the lack of response from S cones in the stimulated region would constrain the inferences downstream stages make about the wavelength composition of small stimuli (Brainard et al., 2008). Second, a recent theory about the neural underpinnings of hue perception posits that only a select group of midget retinal ganglion cells (mRGCs) is involved in color signaling – specifically mRGCs whose L/M centers receive synergistic excitatory input from nearby S cones, rotating their tuning away from the cardinal color axes of the early visual system (Patterson et al., 2019; Schmidt et al., 2014). Cone-to-RGC wiring motifs broadly consistent with this idea have been observed in recent connectomic (Kim et al., 2023, 2024) and physiological (Godat et al., 2024) studies of the primate fovea, although the role these cells play in color perception has not been established.

A prediction common to both theories is that the likelihood of chromatic response would depend on the proximity of S-cones to the stimulated cluster of L and M cones. Specifically, stimuli delivered closer to S cones would be more likely to be seen as chromatic, either because that would facilitate better trichromatic reconstruction (i.e., the non-responsive S cone would rule out a broadband stimulus) or because those L and M cones are more likely to feed into non-canonical cone opponent circuits. To test this prediction, we included the distance to the nearest S cone as a predictor of chromatic response likelihood in our generalized linear mixed-effects model, in addition to L/M heterogeneity and stimulus intensity. Although the latter two factors were significant predictors of chromatic response, we found no significant effect of S cone proximity on how observers categorized the stimuli. We speculate that a more fine-grained characterization of color appearance (i.e., collecting hue scaling responses rather than relying on a red/green/achromatic color naming scheme) could reveal more subtle effects of nearby S cones on color perception (Schmidt, Sabesan, et al., 2018).

## Acknowledgements

The authors thank Austin Roorda and Pavan Tiruveedhula for technical assistance. This work was supported by the Air Force Office of Scientific Research (FA955020-1-0195, FA9550-21-1-0230), National Eye Institute (R01EY023591, R01EY029710, U01EY032055, P30EY003176, P30EY001730, T32EY007043), Alcon Research Institute, Hellman Fellows Program, Burroughs Wellcome Fund Careers at the Scientific Interfaces, and an unrestricted grant from Research to Prevent Blindness.

## Footnotes

1 Caution should be exercised when drawing links between color discrimination and appearance, which can become increasingly decoupled at small spatial scales. In the case of stimuli landing on a spectrally-homogenous patch of receptors, the Principle of Univariance (Rushton, 1972) dictates that discriminating two equiluminant spectral lights should not be possible. However, this does not necessarily imply that said stimuli will appear achromatic. Indeed, minute cone-sized stimuli delivered to single receptors often appear colorful (Hofer, Singer, et al., 2005; Sabesan et al., 2016), even though the individual cells themselves exhibit univariance (Schnapf et al., 1987) and are commonly considered to be “color-blind”.

## References

Arathorn, D. W., Yang, Q., Vogel, C. R., Zhang, Y., Tiruveedhula, P., & Roorda, A. (2007). Retinally stabilized cone-targeted stimulus delivery. Optics Express, 15(21), 13731–13744. 10.1364/OE.15.013731

Asano, Y., Fairchild, M. D., & Blondé, L. (2016). Individual Colorimetric Observer Model. PLOS ONE, 11(2), e0145671-. 10.1371/journal.pone.0145671

Atchison, D. A., & Smith, G. (2005). Chromatic dispersions of the ocular media of human eyes. Journal of the Optical Society of America A, 22(1), 29–37. 10.1364/JOSAA.22.000029

Barbur, J. L., Rodriguez-Carmona, M., Harlow, J. A., Mancuso, K., Neitz, J., & Neitz, M. (2008). A study of unusual Rayleigh matches in deutan deficiency. Visual Neuroscience, 25(3), 507–516. DOI: 10.1017/S0952523808080619

Bieber, M. L., Kraft, J. M., & Werner, J. S. (1998). Effects of known variations in photopigments on L/M cone ratios estimated from luminous efficiency functions. Vision Research, 38(13), 1961–1966. 10.1016/S0042-6989(97)00302-7

Boehm, A. E., Bosten, J., & MacLeod, D. I. A. (2021). Color discrimination in anomalous trichromacy: Experiment and theory. Vision Research, 188, 85–95. 10.1016/j.visres.2021.05.011

Boehm, A. E., MacLeod, D. I. A., & Bosten, J. M. (2014). Compensation for red-green contrast loss in anomalous trichromats. Journal of Vision, 14(13), 19. 10.1167/14.13.19

Boehm, A. E., Privitera, C. M., Schmidt, B. P., & Roorda, A. (2019). Transverse chromatic offsets with pupil displacements in the human eye: sources of variability and methods for real-time correction. Biomedical Optics Express, 10(4), 1691–1706. 10.1364/BOE.10.001691

Bone, R. A., Landrum, J. T., & Cains, A. (1992). Optical density spectra of the macular pigment in vivo and in vitro. Vision Research, 32(1), 105–110. 10.1016/0042-6989(92)90118-3

Bouman, M. A., & Walraven, P. L. (1957). Some Color Naming Experiments for Red and Green Monochromatic Lights. Journal of the Optical Society of America, 47(9), 834–839. 10.1364/JOSA.47.000834

Bowmaker, J. K., Dartnall, H. J., Lythgoe, J. N., & Mollon, J. D. (1978). The visual pigments of rods and cones in the rhesus monkey, Macaca mulatta. The Journal of Physiology, 274(1), 329–348. 10.1113/jphysiol.1978.sp012151

Brainard, D. H. (1997). The Psychophysics Toolbox. Spatial Vision, 10(4), 433–436. 10.1163/156856897X00357

Brainard, D. H., Calderone, J. B., Nugent, A. K., & Jacobs, G. H. (1999). Flicker ERG Responses to Stimuli Parametrically Modulated in Color Space. Investigative Ophthalmology & Visual Science, 40(12), 2840–2847.

Brainard, D. H., Roorda, A., Yamauchi, Y., Calderone, J. B., Metha, A., Neitz, M., Neitz, J., Williams, D. R., & Jacobs, G. H. (2000). Functional consequences of the relative numbers of L and M cones. Journal of the Optical Society of America A, 17(3), 607–614. 10.1364/josaa.17.001684

Brainard, D. H., Williams, D. R., & Hofer, H. (2008). Trichromatic reconstruction from the interleaved cone mosaic: Bayesian model and the color appearance of small spots. Journal of Vision, 8(5), 15.1-23. 10.1167/8.5.15

Carroll, J., Neitz, J., & Neitz, M. (2002). Estimates of L: M cone ratio from ERG flicker photometry and genetics. Journal of Vision, 2(8), 1. 10.1167/2.8.1

Chaparro, A., Stromeyer, C. F., Kronauer, R. E., & Eskew, R. T. (1994). Separable red-green and luminance detectors for small flashes. Vision Research, 34(6), 751–762. 10.1016/0042-6989(94)90214-3

Cicerone, C. M., & Nerger, J. L. (1989). The relative numbers of long-wavelength-sensitive to middle-wavelength-sensitive cones in the human fovea centralis. Vision Research, 29(1), 115–128. 10.1016/0042-6989(89)90178-8

CIE. (2006). CIE 2006 LMS cone fundamentals for 2° field size in terms of energy. 10.25039/CIE.DS.tijidesg

Danilova, M. V, Chan, C. H., & Mollon, J. D. (2013). Can spatial resolution reveal individual differences in the L:M cone ratio? Vision Research, 78, 26–38. 10.1016/j.visres.2012.12.006

de Vries, H. L. (1946). Luminosity Curve of Trichromats. Nature, 157(3996), 736–737. 10.1038/157736b0

de Vries, H. L. (1949). The heredity of the relative numbers of red and green receptors in the human eye. Genetica, 24(1), 199–212. 10.1007/BF01487206

Emery, K. J., Isherwood, Z. J., & Webster, M. A. (2023). Gaining the system: limits to compensating color deficiencies through post-receptoral gain changes. Journal of the Optical Society of America A, 40(3), A16–A25. 10.1364/JOSAA.480035

Field, G. D., Gauthier, J. L., Sher, A., Greschner, M., Machado, T. a., Jepson, L. H., Shlens, J., Gunning, D. E., Mathieson, K., Dabrowski, W., Paninski, L., Litke, A. M., & Chichilnisky, E. J. (2010). Functional connectivity in the retina at the resolution of photoreceptors. Nature, 467(7316), 673–677. 10.1038/nature09424

Finkelstein, M. A., & Hood, D. C. (1984). Detection and discrimination of small, brief lights: Variable tuning of opponent channels. Vision Research, 24(3), 175–181. 10.1016/0042-6989(84)90119-6

Fong, J., Doyle, H. K., Wang, C., Boehm, A. E., Herbeck, S. R., Pandiyan, V. P., Schmidt, B. P., Tiruveedhula, P., Vanston, J. E., Tuten, W. S., Sabesan, R., Roorda, A., & Ng, R. (2025). Novel color via stimulation of individual photoreceptors at population scale. Science Advances, 11(16), eadu1052. 10.1126/sciadv.adu1052

Garrigan, P., Ratliff, C. P., Klein, J. M., Sterling, P., Brainard, D. H., & Balasubramanian, V. (2010). Design of a Trichromatic Cone Array. PLOS Computational Biology, 6(2), e1000677-. 10.1371/journal.pcbi.1000677

Godat, T., Kohout, K., Parkins, K., Yang, Q., McGregor, J. E., Merigan, W. H., Williams, D. R., & Patterson, S. S. (2024). Cone-Opponent Ganglion Cells in the Primate Fovea Tuned to Noncardinal Color Directions. Journal of Neuroscience, 44(18). 10.1523/JNEUROSCI.1738-23.2024

Greene, M. J., Boehm, A. E., Vanston, J. E., Pandiyan, V. P., Sabesan, R., & Tuten, W. S. (2024). Unique yellow shifts for small and brief stimuli in the central retina. Journal of Vision, 24(6), 2. 10.1167/jov.24.6.2

Gunther, K. L., & Dobkins, K. R. (2002). Individual differences in chromatic (red/green) contrast sensitivity are constrained by the relative number of L-versus M-cones in the eye. Vision Research, 42(11), 1367–1378. 10.1016/S0042-6989(02)00043-3

Hagstrom, S. A., Neitz, J., & Neitz, M. (1998). Variations in cone populations for red–green color vision examined by analysis of mRNA. NeuroReport, 9(9). 10.1097/00001756-199806220-00009

Hagstrom, S. A., Neitz, M., & Neitz, J. (2000). Cone pigment gene expression in individual photoreceptors and the chromatic topography of the retina. Journal of the Optical Society of America A, 17(3), 527–537. 10.1364/JOSAA.17.000527

Harmening, W. M., Tiruveedhula, P., Roorda, A., & Sincich, L. C. (2012). Measurement and correction of transverse chromatic offsets for multi-wavelength retinal microscopy in the living eye. Biomedical Optics Express, 3(9), 2066–2077. 10.1364/BOE.3.002066

Harmening, W. M., Tuten, W. S., Roorda, A., & Sincich, L. C. (2014). Mapping the perceptual grain of the human retina. Journal of Neuroscience, 34(16). 10.1523/JNEUROSCI.5191-13.2014

He, J., Taveras-Cruz, Y., & Eskew Jr., R. T. (2021). Modeling individual variations in equiluminance settings. Journal of Vision, 21(7), 15. 10.1167/jov.21.7.15

Hofer, H., Carroll, J., Neitz, J., Neitz, M., & Williams, D. R. (2005). Organization of the human trichromatic cone mosaic. Journal of Neuroscience, 25(42), 9669–9679. 10.1523/JNEUROSCI.2414-05.2005

Hofer, H., Singer, B., & Williams, D. R. (2005). Different sensations from cones with the same photopigment. Journal of Vision, 5(5), 444–454. 10:1167/5.5.5

Hood, S. M., Mollon, J. D., Purves, L., & Jordan, G. (2006). Color discrimination in carriers of color deficiency. Vision Research, 46(18), 2894–2900. 10.1016/j.visres.2006.02.028

Jordan, G., & Mollon, J. D. (1997). Unique hues in heterozygotes for protan and deutan deficiencies. *Colour Vision Deficiencies XIII: Proceedings of the Thirteenth Symposium of the International Research Group on Colour Vision Deficiencies, Held in Pau*, France July *27–30*, 1995, 67–76.

Kim, Y. J., Packer, O., & Dacey, D. M. (2024). A circuit motif for color in the human foveal retina. Proceedings of the National Academy of Sciences of the United States of America, 121(36). 10.1073/pnas.2405138121

Kim, Y. J., Packer, O., Pollreisz, A., Martin, P. R., Grünert, U., & Dacey, D. M. (2023). Comparative connectomics reveals noncanonical wiring for color vision in human foveal retina. Proceedings of the National Academy of Sciences of the United States of America, 120(118). 10.1073/pnas.2300545120

King-Smith, P. E., & Carden, D. (1976). Luminance and opponent-color contributions to visual detection and adaptation and to temporal and spatial integration. Journal of the Optical Society of America, 66(7), 709–717. 10.1364/JOSA.66.000709

Knoblauch, K., Marsh-Armstrong, B., & Werner, J. S. (2020). Suprathreshold contrast response in normal and anomalous trichromats. Journal of the Optical Society of America A, 37(4), A133–A144. 10.1364/JOSAA.380088

Kraft, T. W., Neitz, J., & Neitz, M. (1998). Spectra of human L cones. Vision Research, 38(23), 3663–3670. 10.1016/S0042-6989(97)00371-4

Krauskopf, J. (1964). Color Appearance of Small Stimuli and the Spatial Distribution of Color Receptors. Journal of the Optical Society of America, 54(9), 1171. 10.1364/JOSA.54.001171

Krauskopf, J. (2000). Relative number of long- and middle-wavelength-sensitive cones in the human fovea. Journal of the Optical Society of America A, 17(3), 510–516. 10.1364/JOSAA.17.000510

Krauskopf, J., & Srebro, R. (1965). Spectral Sensitivity of Color Mechanisms: Derivation from Fluctuations of Color Appearance near Threshold. Science, 150(3702), 1477–1479. 10.1126/science.150.3702.1477

Kremers, J., Scholl, H. P. N., Knau, H., Berendschot, T. T. J. M., Usui, T., & Sharpe, L. T. (2000). L/M cone ratios in human trichromats assessed by psychophysics, electroretinography, and retinal densitometry. Journal of the Optical Society of America A, 17(3), 517–526. 10.1364/JOSAA.17.000517

Lee, B. B., Pokorny, J., Smith, V. C., Martin, P. R., & Valbergt, A. (1990). Luminance and chromatic modulation sensitivity of macaque ganglion cells and human observers. Journal of the Optical Society of America A, 7(12), 2223–2236. 10.1364/JOSAA.7.002223

Li, P. H., Field, G. D., Greschner, M., Ahn, D., Gunning, D. E., Mathieson, K., Sher, A., Litke, A. M., & Chichilnisky, E. J. (2014). Retinal representation of the elementary visual signal. Neuron, 81(1), 130–139. 10.1016/j.neuron.2013.10.043

MacLeod, D. I. A., Williams, D. R., & Makous, W. (1992). A visual nonlinearity fed by single cones. Vision Research, 32(2), 347–363. 10.1016/0042-6989(92)90144-8

MacLeod, D. I., & Boynton, R. M. (1979). Chromaticity diagram showing cone excitation by stimuli of equal luminance. Journal of the Optical Society of America, 69(8), 1183–1186. 10.1364/JOSA.69.001183

Manning, J. R., & Brainard, D. H. (2009). Optimal design of photoreceptor mosaics: Why we do not see color at night. Visual Neuroscience, 26(1), 5–19. DOI: 10.1017/S095252380808084X

Miyahara, E., Pokorny, J., Smith, V. C., Baron, R., & Baron, E. (1998). Color vision in two observers with highly biased LWS/MWS cone ratios. Vision Research, 38(4), 601–612. 10.1016/S0042-6989(97)88334-4

Mollon, J. D. (2006). Monge: The Verriest Lecture, Lyon, July 2005. *Visual Neuroscience*, *23*(3–4), 297–309. DOI: 10.1017/S0952523806233479

Mollon, J. D., & Bowmaker, J. K. (1992). The spatial arrangement of cones in the primate fovea. Nature, 360(6405), 677–679. 10.1038/360677a0

Mozaffari, S., LaRocca, F., Jaedicke, V., Tiruveedhula, P., & Roorda, A. (2020). Wide-vergence, multi-spectral adaptive optics scanning laser ophthalmoscope with diffraction-limited illumination and collection. Biomedical Optics Express, 11(3), 1617–1632. 10.1364/BOE.384229

Mullen, K. T. (1985). The contrast sensitivity of human colour vision to red-green and blue-yellow chromatic gratings. The Journal of Physiology, 359(1), 381–400. 10.1113/jphysiol.1985.sp015591

Neitz, A., Jiang, X., Kuchenbecker, J. A., Domdei, N., Harmening, W., Yan, H., Yeonan-Kim, J., Patterson, S. S., Neitz, M., Neitz, J., Coates, D. R., & Sabesan, R. (2020). Effect of cone spectral topography on chromatic detection sensitivity. Journal of the Optical Society of America A, 37(4), A244–A254. 10.1364/JOSAA.382384

Neitz, J., Carroll, J., Yamauchi, Y., Neitz, M., & Williams, D. R. (2002). Color perception is mediated by a plastic neural mechanism that is adjustable in adults. Neuron, 35(4), 783– 792. 10.1016/S0896-6273(02)00818-8

Neitz, J., & Jacobs, G. H. (1986). Polymorphism of the long-wavelength cone in normal human colour vision. Nature, 323(6089), 623–625. 10.1038/323623a0

Neitz, J., & Neitz, M. (2011). The genetics of normal and defective color vision. Vision Research, 51(7), 633–651. 10.1016/j.visres.2010.12.002

Otake, S., Gowdy, P. D., & Cicerone, C. M. (2000). The spatial arrangement of L and M cones in the peripheral human retina. Vision Research, 40(6), 677–693. 10.1016/S0042-6989(99)00202-3

Pandiyan, V. P., Maloney-Bertelli, A., Kuchenbecker, J. A., Boyle, K. C., Ling, T., Chen, Z. C., Park, B. H., Roorda, A., Palanker, D., & Sabesan, R. (2020). The optoretinogram reveals the primary steps of phototransduction in the living human eye. Science Advances, 6(37), eabc1124. 10.1126/sciadv.abc1124

Pandiyan, V. P., Schleufer, S., Slezak, E., Fong, J., Upadhyay, R., Roorda, A., Ng, R., & Sabesan, R. (2022). Characterizing cone spectral classification by optoretinography. Biomedical Optics Express, 13(12), 6574–6594. 10.1364/BOE.473608

Patterson, S. S., Neitz, M., & Neitz, J. (2019). Reconciling Color Vision Models With Midget Ganglion Cell Receptive Fields. In Frontiers in Neuroscience (Vol. 13). Frontiers Media S.A. 10.3389/fnins.2019.00865

Roorda, A., Metha, A. B., Lennie, P., & Williams, D. R. (2001). Packing arrangement of the three cone classes in primate retina. Vision Research, 41(10), 1291–1306. 10.1016/S0042-6989(01)00043-8

Roorda, A., Romero-Borja, F., Donnelly III, W. J., Queener, H., Hebert, T. J., & Campbell, M. C. W. (2002). Adaptive optics scanning laser ophthalmoscopy. Optics Express, 10(9), 405–412. 10.1364/OE.10.000405

Roorda, A., & Williams, D. R. (1999). The arrangement of the three cone classes in the living human eye. Nature, 397(6719), 520–522. 10.1038/17383

Rushton, W. A. H. (1972). Review lecture. Pigments and signals in colour vision. The Journal of Physiology, 220(3), 1–31. 10.1113/jphysiol.1972.sp009719

Rushton, W. A. H., & Baker, H. D. (1964). Red/green sensitivity in normal vision. Vision Research, 4(1), 75–85. 10.1016/0042-6989(64)90034-3

Rushton, W. A. H., & Henry, G. H. (1968). Bleaching and regeneration of cone pigments in man. Vision Research, 8(6), 617–631. 10.1016/0042-6989(68)90040-0

Sabesan, R., Schmidt, B. P., Tuten, W. S., & Roorda, A. (2016). The elementary representation of spatial and color vision in the human retina. Science Advances, 2(9). 10.1126/sciadv.1600797

Schmidt, B. P., Boehm, A. E., Foote, K. G., & Roorda, A. (2018). The spectral identity of foveal cones is preserved in hue perception. Journal of Vision, 18(11), 19. 10.1167/18.11.19

Schmidt, B. P., Boehm, A. E., Tuten, W. S., & Roorda, A. (2019). Spatial summation of individual cones in human color vision. PLoS ONE, 14(7). 10.1371/journal.pone.0211397

Schmidt, B. P., Neitz, M., & Neitz, J. (2014). Neurobiological hypothesis of color appearance and hue perception. JOSA A, 31(4), A195–A207. 10.1364/JOSAA.31.00A195

Schmidt, B. P., Sabesan, R., Tuten, W. S., Neitz, J., & Roorda, A. (2018). Sensations from a single M-cone depend on the activity of surrounding S-cones. Scientific Reports, 8(1), 1–10. 10.1038/s41598-018-26754-1

Schnapf, J. L., Kraft, T. W., & Baylor, D. A. (1987). Spectral sensitivity of human cone photoreceptors. Nature, 325(6103), 439–441. 10.1038/325439a0

Sekiguchi, N., Williams, D. R., & Brainard, D. H. (1993). Efficiency in detection of isoluminant and isochromatic interference fringes. Journal of the Optical Society of America A, 10(10), 2118–2133. http://www.ncbi.nlm.nih.gov/pubmed/8229351

Shi, K., Luo, M. R., Rider, A. T., Huang, T., Xu, L., & Stockman, A. (2024). A multi-primary trichromator to derive individual color matching functions and cone spectral sensitivities. Color Research & Application, 49(5), 449–464. 10.1002/col.22928

Stockman, A., Jägle, H., Pirzer, M., & Sharpe, L. T. (2008). The dependence of luminous efficiency on chromatic adaptation. Journal of Vision, 8(16), 1. 10.1167/8.16.1

Stockman, A., Sharpe, L. T., & Fach, C. (1999). The spectral sensitivity of the human short-wavelength sensitive cones derived from thresholds and color matches. Vision Research, 39(17), 2901–2927. 10.1016/S0042-6989(98)00225-9

Thomas, P. B. M., & Mollon, J. D. (2004). Modelling the Rayleigh match. Visual Neuroscience, 21(3), 477–482. DOI: 10.1017/S095252380421344X

Tregillus, K. E. M., Isherwood, Z. J., Vanston, J. E., Engel, S. A., MacLeod, D. I. A., Kuriki, I., & Webster, M. A. (2021). Color Compensation in Anomalous Trichromats Assessed with fMRI. Current Biology, 31(5), 936–942.e4. 10.1016/j.cub.2020.11.039

Vimal, R. L. P., Pokorny, J., Smith, V. C., & Shevell, S. K. (1989). Foveal cone thresholds. Vision Research, 29(1), 61–78. 10.1016/0042-6989(89)90174-0

Vorobyev, M., & Osorio, D. (1998). Receptor noise as a determinant of colour thresholds. Proceedings of the Royal Society of London. Series B: Biological Sciences, 265(1394), 351–358. 10.1098/rspb.1998.0302

Watson, A. B. (2017). QUEST+: A general multidimensional Bayesian adaptive psychometric method. Journal of Vision, 17(3), 10. 10.1167/17.3.10

Welbourne, L. E., Morland, A. B., & Wade, A. R. (2015). Human colour perception changes between seasons. Current Biology, 25(15), R646–R647. 10.1016/j.cub.2015.06.030

Williams, D. R., MacLeod, D. I. A., & Hayhoe, M. M. (1981). Punctate sensitivity of the blue-sensitive mechanism. Vision Research, 21(9), 1357–1375. 10.1016/0042-6989(81)90242-X

Wuerger, S., Ashraf, M., Kim, M., Martinovic, J., Pérez-Ortiz, M., & Mantiuk, R. K. (2020). Spatio-chromatic contrast sensitivity under mesopic and photopic light levels. Journal of Vision, 20(4), 23. 10.1167/jov.20.4.23

Yang, Q., Arathorn, D. W., Tiruveedhula, P., Vogel, C. R., & Roorda, A. (2010). Design of an integrated hardware interface for AOSLO image capture and cone-targeted stimulus delivery. Optics Express, 18(17), 17841–17858. 10.1364/OE.18.017841

Zhang, F., Kurokawa, K., Lassoued, A., Crowell, J. A., & Miller, D. T. (2019). Cone photoreceptor classification in the living human eye from photostimulation-induced phase dynamics. Proceedings of the National Academy of Sciences, 116(16), 7951–7956. 10.1073/pnas.1816360116

Zhang, L. Q., Cottaris, N. P., & Brainard, D. H. (2022). An image reconstruction framework for characterizing initial visual encoding. ELife, 11, 1–30. 10.7554/eLife.71132

Zhaoping, L., & Carroll, J. (2016). An analytical model of the influence of cone sensitivity and numerosity on the Rayleigh match. Journal of the Optical Society of America A, 33(3), A228–A237. 10.1364/JOSAA.33.00A228

